## Supplemental material for "The nuts and bolts of SARS-CoV-2 Spike Receptor Binding Domain heterologous expression"

### SUPPLEMENTARY MATERIAL

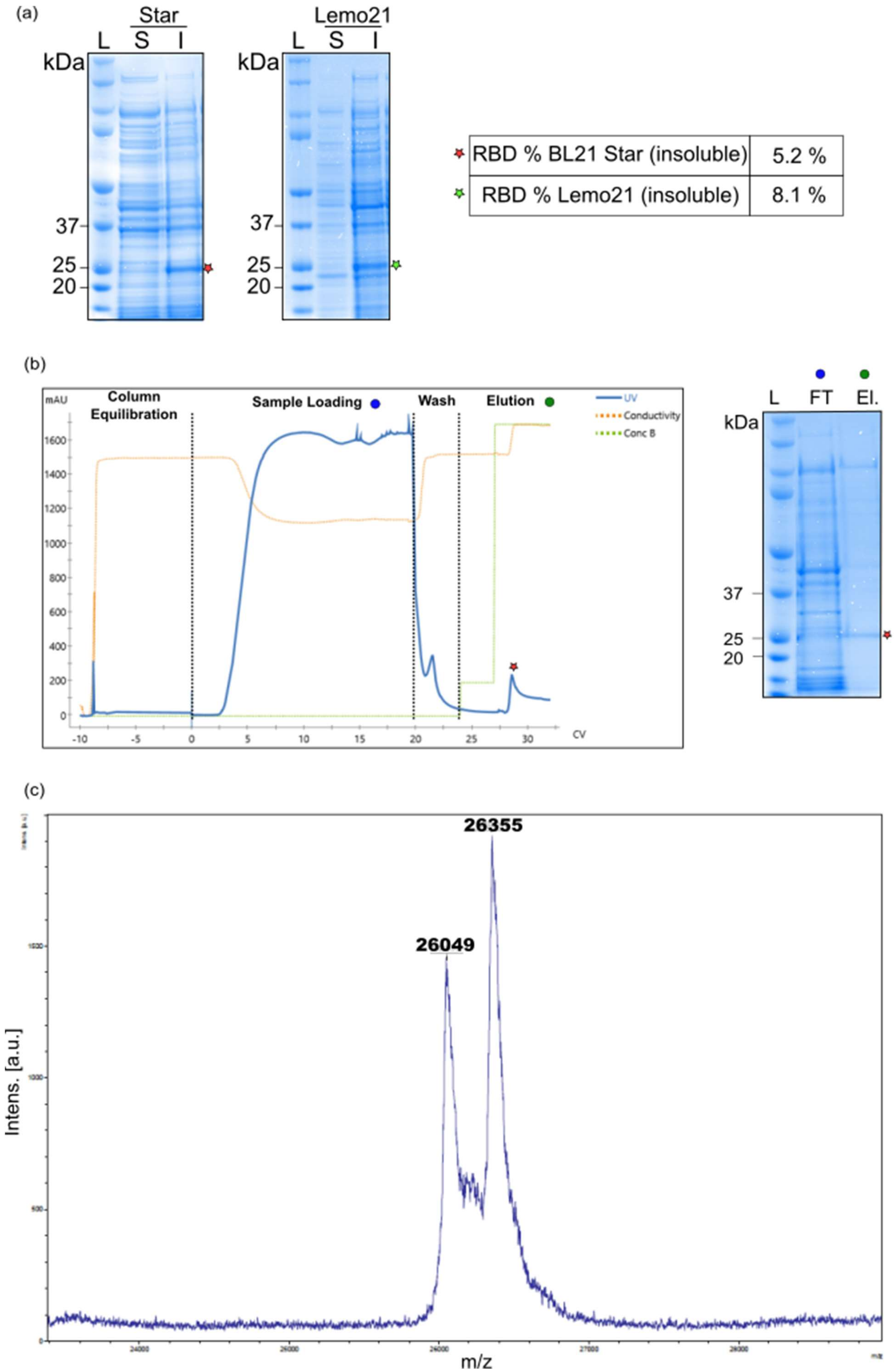
(a)
Star
L S I
kDa
37
25
20
Lemo21
L S I
kDa
37
25
20
RBD % BL21 Star (insoluble) 5.2 %
RBD % Lemo21 (insoluble) 8.1 %
(b)
mAU
Column Equilibration
Sample Loading
Wash
Elution
UV
Conductivity
Conc B
CV
L FT El.
kDa
37
25
20
(c)
26355
26049
Intens. [a.u.]
m/z

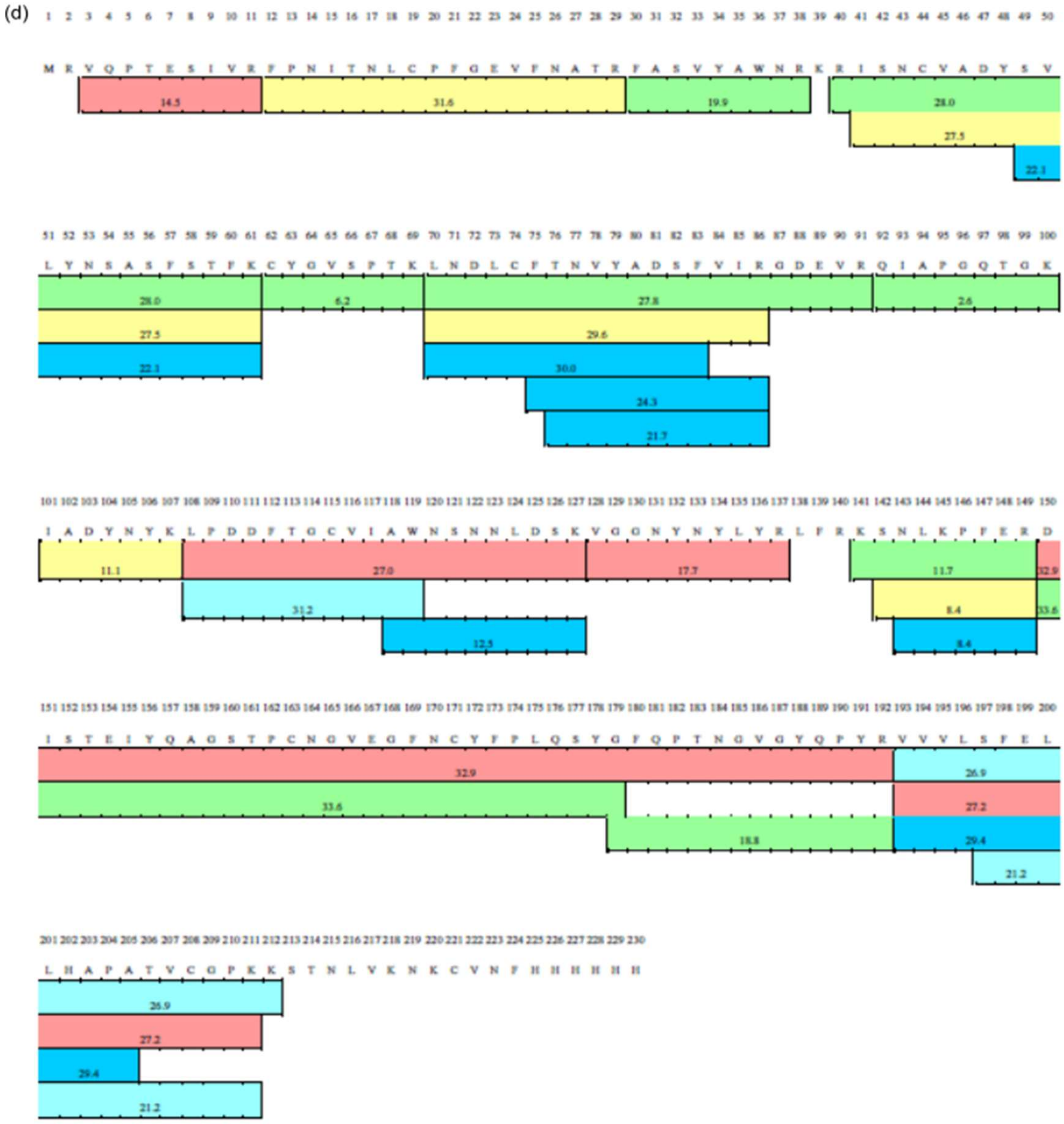

**Supplementary Figure 1.** (a) SDS-PAGE analysis of the soluble (S) and insoluble fractions (I) from RBD expression in BL21 Star strain (left) and Lemo21 (right) L= molecular weight ladder. (b) Chromatogram profile of IMAC purification under denaturing conditions of RBD protein (left) and corresponding SDS-PAGE (4-12%) analysis of flow through (FT) and elution (El) fractions. (c) MALDI mass spectrum of RBD sample produced in *E. coli*. (d) Schematic view of recombinant RBD sequence coverage. Protein of interest was represented by its aminoacidic sequence in FASTA and a colored frame in correspondence to each peptide experimentally identified was reported. The rainbow color of the frame is related to the signal intensity, the red one being the most intense and the blue one the lowest. The peptide mass fingerprint (PMF) was determined considering both the molecular weight (from the full mass) and the fragmentation (from the MS/MS mass spectra) of all the peptides detected in the LC-MS/MS analysis of each sample. The result obtained, expressed as sequence coverage (%) for the recombinant *E. coli*-RBD protein, was 89.6%.

(a)

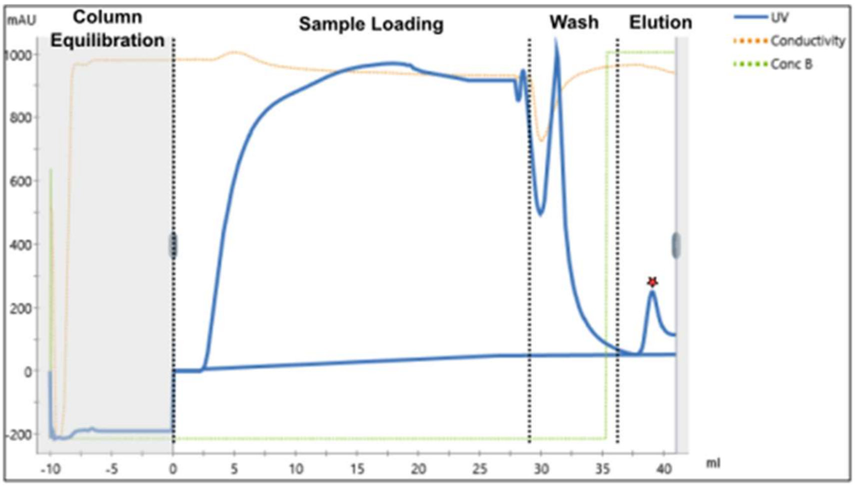
mAU
Column
Equilibration
Sample Loading
Wash
Elution
UV
Conductivity
Conc B
1000
800
600
400
200
0
-200
-10
-5
0
5
10
15
20
25
30
35
40
ml

(b)

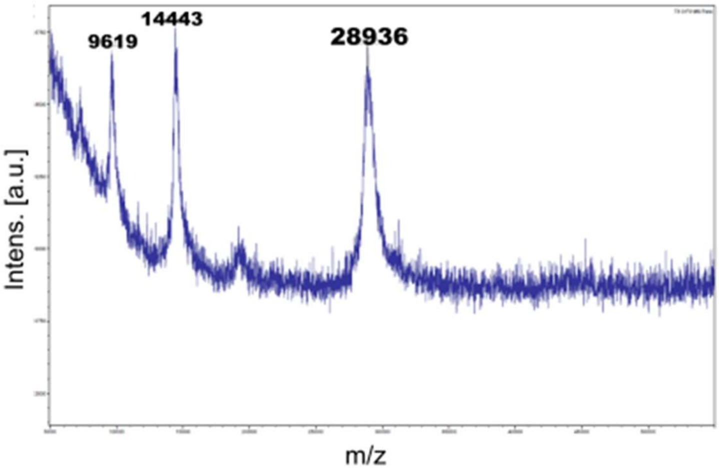
9619
14443
28936
Intens. [a.u.]
m/z

(c)

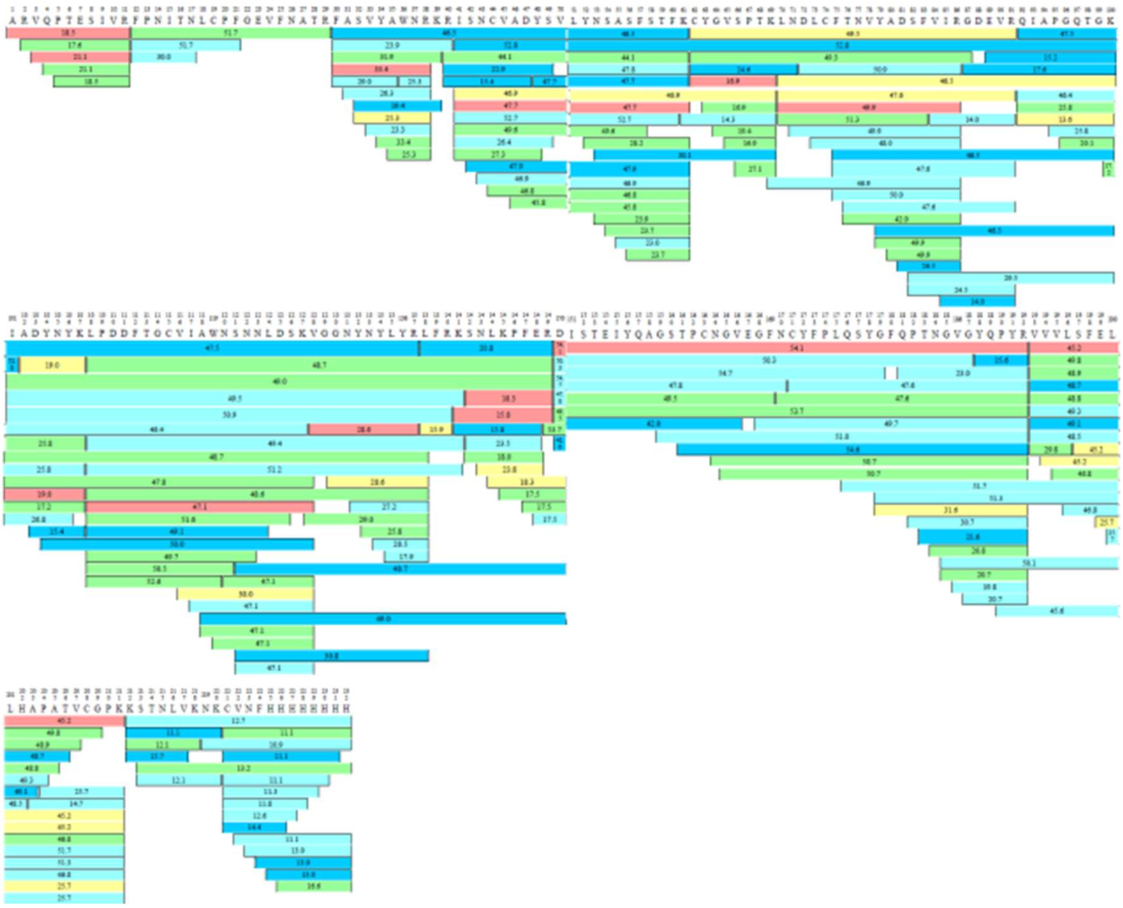

(d)

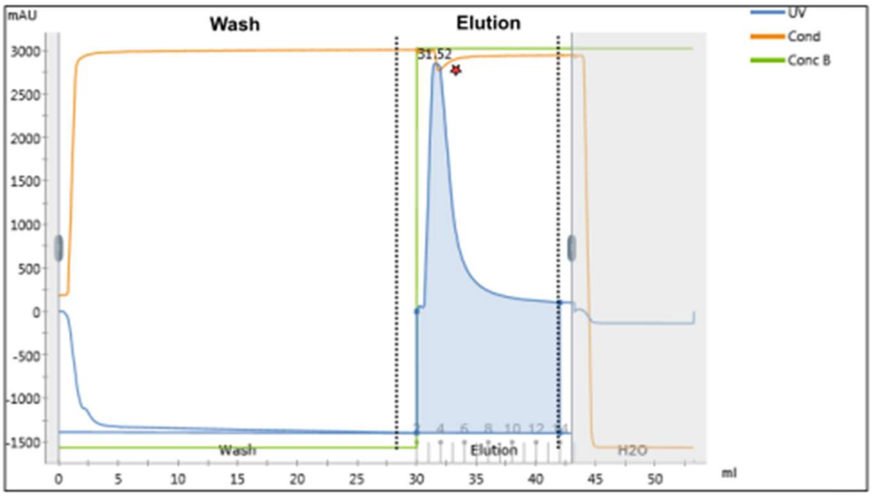
mAU
Wash
Elution
UV
Cond
Conc B
31.52
3000
2500
2000
1500
1000
500
0
-500
-1000
-1500
0
5
10
15
20
25
30
35
40
45
50
ml
Wash
Elution
H2O

(e)

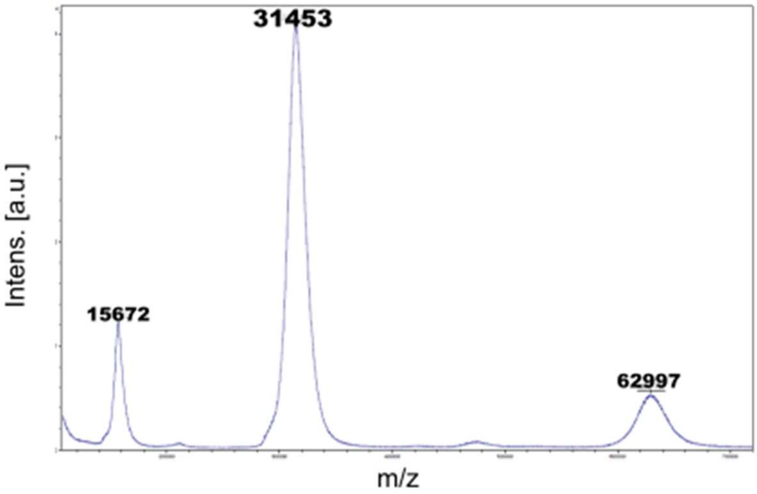
31453
15672
62997
Intens. [a.u.]
m/z

(f)

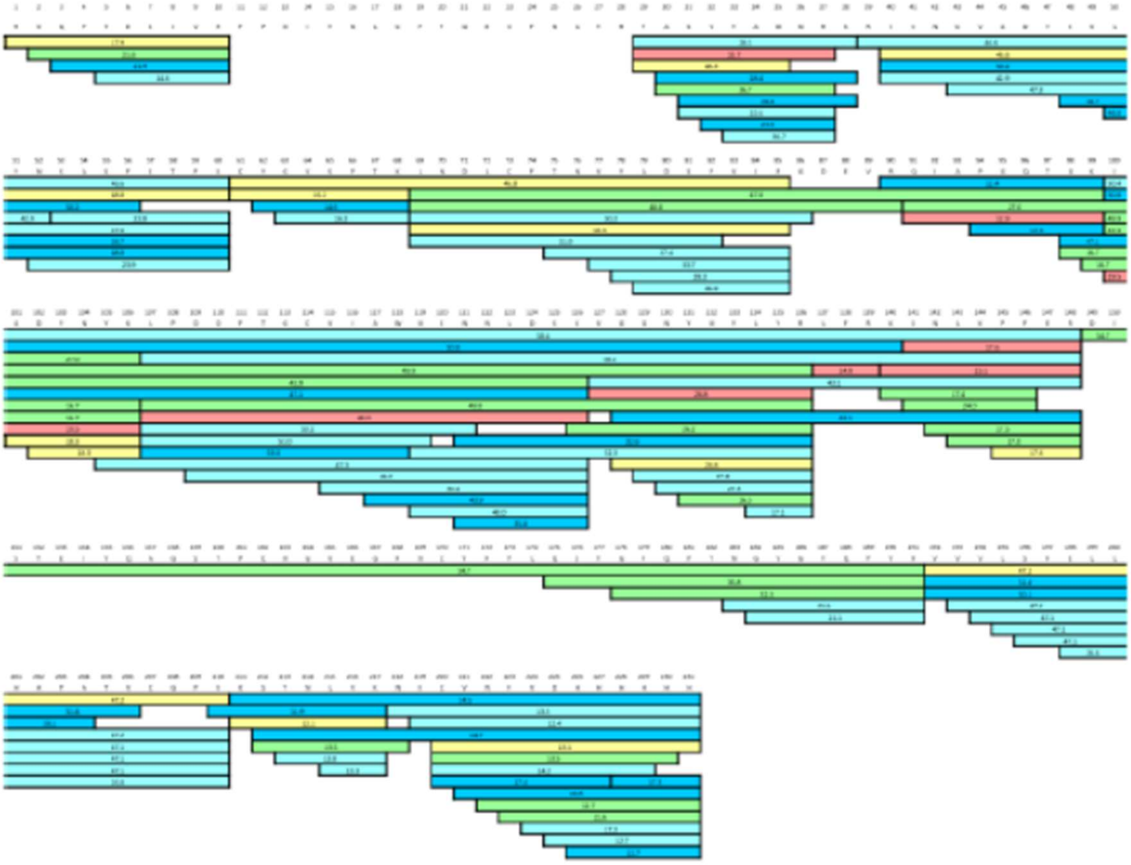

**Supplementary Figure 2.** (a) Chromatogram profile of IMAC purification of RBD protein secreted from Hi-5 cells. (b) MALDI mass spectrum of RBD sample produced in insect cells. The main peak (m/z 28936) corresponds to glycosylated RBD, while the peaks at m/z 14433 and 9619 are attributable to doubly and triply charged species of the same protein. (c) Schematic view of recombinant insect-RBD sequence coverage. The peptide mass fingerprint (PMF) was determined considering both the molecular weight (from the full mass) and the fragmentation (from the MS/MS mass spectra) of all the peptides detected in the LC-MS/MS analysis of each sample. The result obtained, expressed as sequence coverage (%) for the recombinant RBD protein (insect), was 100%. (d) Chromatogram profile of the IMAC purification of RBD protein secreted from mammalian HEK-293. (e) MALDI mass spectrum of RBD sample produced in HEK-293. The mass spectrum showed a main peak at m/z 31453 (glycosylated RBD), a doubly charge species at m/z 15672 and a dimeric species (formed inside the mass spectrometer) at m/z 62997. (f) Schematic view of recombinant HEK-293-RBD sequence coverage. The peptide mass fingerprint (PMF) was determined considering both the molecular weight (from the full mass) and the fragmentation (from the MS/MS mass spectra) of all the peptides detected in the LC-MS/MS analysis of each sample. The result obtained, expressed as sequence coverage (%) for the recombinant RBD protein (insect), was 92.2%. The protein(s) was represented by its aminoacidic sequence in FASTA and a colored frame in correspondence to each peptide experimentally identified was reported. The rainbow color of the frame indicates signal intensity (red is the most intense signal, blue is the lowest one).

(a)

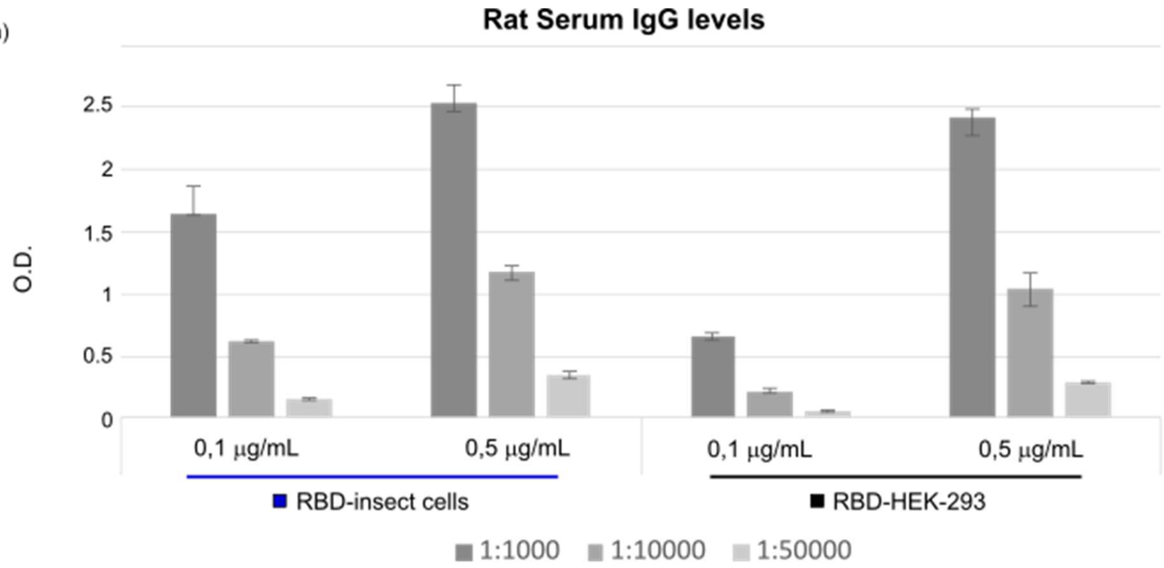


(b)

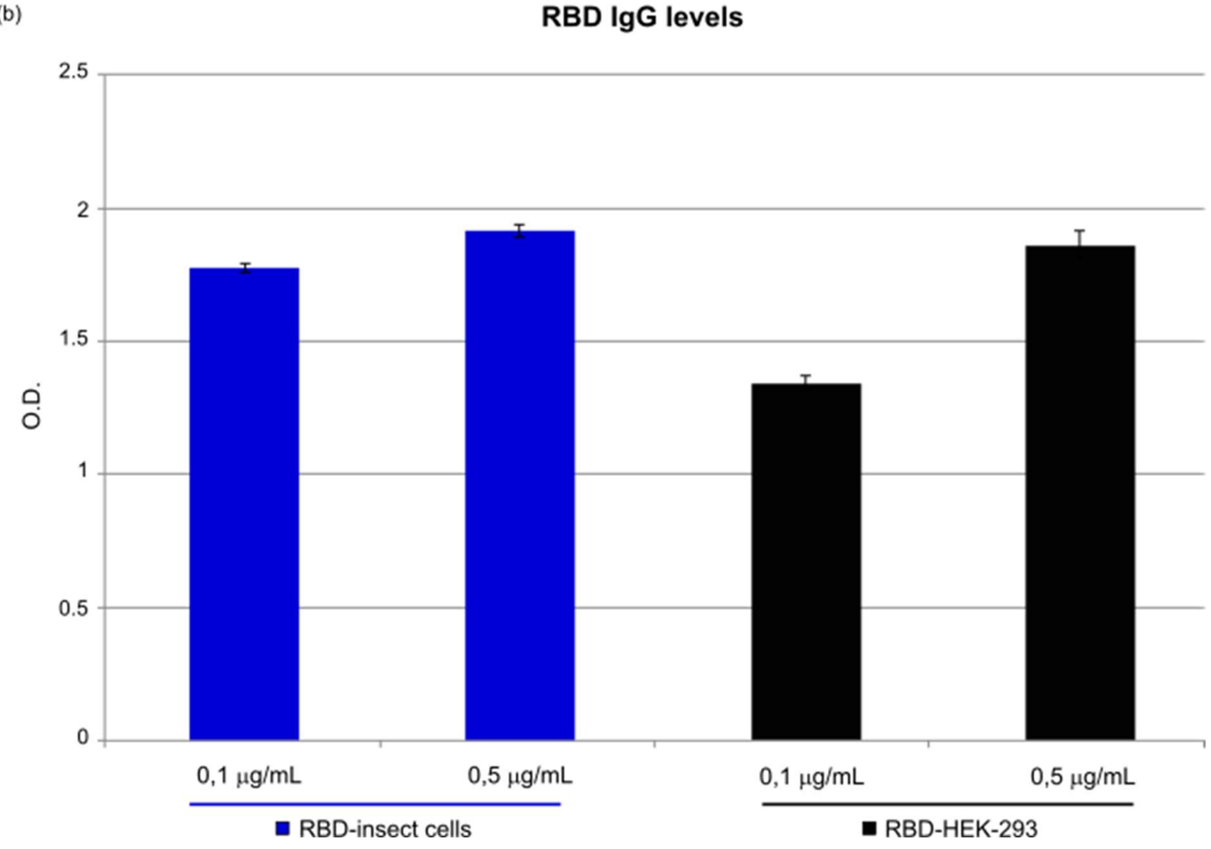


**Supplementary Figure 3.** (a) Serum from immunized rat against Covid-19 was used to compare different concentrations (0.1 and 0.5 μg/mL) of RBD produced in insect (blue) and HEK-293 cells (black). Optical density (OD) was measured at 405 nm and it is represented in the y-axis. The x-axis shows the dilution factor for serum samples (1:1000, 1:10000, 1:50000). Bars indicate standard deviations. (b) Commercial antibody against the S1 subunit of SARS-CoV-2 Spike was used to compare different concentrations (0.1 and 0.5 μg/mL) of

RBD produced in insect (blue) and HEK-293 cells (black). Optical density (OD) was measured at 450 nm and bars indicate standard deviations.

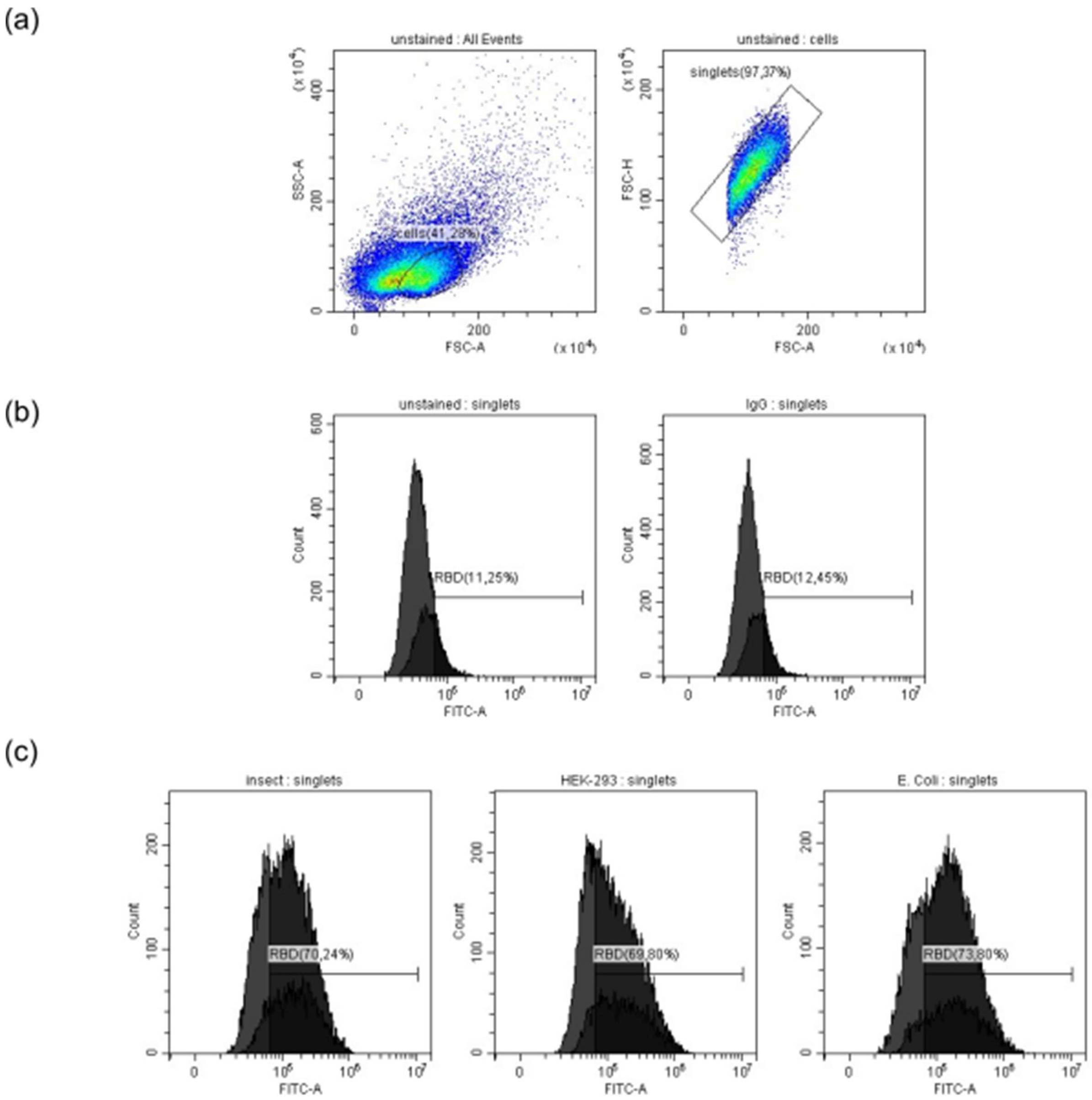


**Supplementary Figure 4.** Representative flow cytometry plots showing (a) gating strategy: dead cells were excluded by their distinctive smaller size (FSC-A) and single cells gated using FSC-H/FSC-A gating. (b) Vero E6 cells alone (left) or incubated only with the secondary antibody (right). (c) Vero E6 cells incubated with insect-RBD (left panel), HEK-293-RBD (middle panel) and *E.* coli-RBD (right panel) followed by anti-RBD primary antibody and secondary antibody.

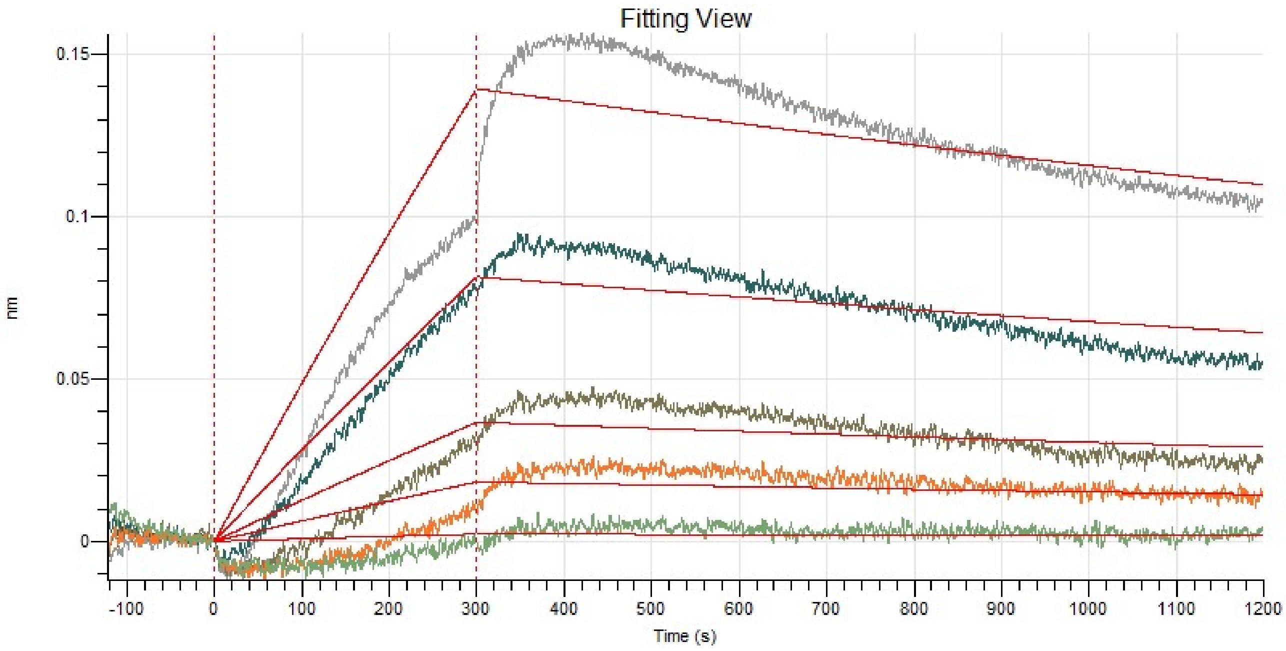


**Supplementary Figure 5.** BLI data for the binding of *E.* coli-RBD to ACE2-Fc. After a baseline step, sensorgram starts with the association (0 - 300 s) of RBD on the ACE2 sensor, followed by the dissociation phase (900 s).
